# *FLOWERING LOCUS T1* is a pleiotropic regulator of reproductive development, plant longevity, and source-sink relations in barley

**DOI:** 10.1101/2025.11.07.687273

**Authors:** Gesa Helmsorig, Tianyu Lan, Einar B. Haraldsson, Thea Rütjes, Philipp Westhoff, Katrin Weber, Jochen Kumlehn, Götz Hensel, Rüdiger Simon, Maria von Korff

**Author notes:** contributed equally (Gesa Helmsorig, Tianyu Lan).

## Abstract

Source–sink interactions play a critical but mechanistically underexplored role in coordinating reproductive output and longevity in plants. Here, we investigated the role of *FT1*, the barley homolog of the florigen *FLOWERING LOCUS T* (*FT*), in regulating source (leaf) and sink (inflorescence) development and metabolism. Using *ft1* knock-out mutants in the spring barley cultivar Golden Promise, which carries a mutated *ppd-H1* allele, and in an introgression line with a wild-type *Ppd-H1* allele, we show that *Ppd-H1* primarily regulates the timing of inflorescence development and flowering through *FT1*, whereas variation in tiller number and leaf size is determined by the genetic background. *ft1* mutants exhibited reduced determinacy of both leaf and inflorescence meristems, resulting in increased leaf and spikelet numbers and size, but severely reduced inflorescence fertility, altered senescence patterns, and significantly extended plant longevity. The *ft1* mutants exhibited a strong transcriptional reprogramming of genes involved in both the light and dark reactions of photosynthesis in the leaf, alongside an upregulation of genes associated with carbon catabolism and stress responses in the leaf and inflorescence.

Elevated soluble sugar and starch levels in ft1 inflorescences indicated that the impaired development and fertility of *ft1* inflorescences were not caused by carbon limitation, but instead reflected a reduced sink strength. Our work reveals that *FT1* coordinates the development of vegetative and reproductive meristems and organs with plant physiology and metabolism, thereby regulating source–sink relationships and balancing plant longevity with reproductive output.

## Introduction

The timing of flowering is a critical determinant of both plant reproductive success and crop yield. Precise coordination of flowering time with environmental conditions is essential to maximise seed set and ensure successful reproduction. At the same time, the developmental timing strongly influences overall shoot and inflorescence architecture and resource allocations, which in turn impact yield potential (Gol et al. 2017, Bi et al. 2019, Shanmugaraj et al. 2023, Dresselhaus et al. 2025). The above-ground architecture of a plant is mainly defined by the number, size, and arrangement of organs formed from the shoot apical meristem (SAM) (Benlloch et al., 2007). The number of organs developed throughout the lifecycle of a plant depends on the identity and determinacy of shoot meristems. During vegetative growth, the SAM produces leaf primordia and axillary meristems (AXMs) that grow out into leaves and branches, respectively (Wang and Smith, 2018). Reproductive growth is initiated by the irreversible transition of a vegetative SAM into a reproductive inflorescence meristem (IM), which ceases to produce leaf primordia and instead forms floral meristems in eudicots and spikelet meristems (SMs) in grasses (Bommert and Whipple, 2018). Meristem determinacy controls how long a meristem remains active and thus the number of organs developed by each meristem type (Bartlett and Thompson, 2014). A major gene affecting meristem identity and determinacy in *Arabidopsis thaliana* is *FLOWERING LOCUS T (FT),* which was identified as the florigen, a mobile floral stimulus that transfers the information to induce flowering from leaves to the developing shoot apex (Chailakhyan, 1936; Corbesier et al., 2007; Tamaki et al., 2007). In flowering-inducing conditions, *FT* is upregulated in phloem companion cells in the vascular tissue of the leaf and transported as a protein through the phloem to the shoot apex (Kardailsky et al., 1999; Corbesier et al., 2007; Tamaki et al., 2007; Chen et a., 2018). In the shoot apex, the FT protein interacts with FLOWERING LOCUS D (FD), bridged by 14-3-3 proteins, to form the florigen activation complex (FAC) and activate the expression of floral meristem identity genes and promote floral organ differentiation (Abe et al., 2005; Wigge et al., 2005; Taoka et al., 2011). Through its effects on meristem identity and determinacy, FT controls the length of the vegetative (source-building) phase and thus the number of leaves and potential branches, as well as sink potential, the number of flowers and the size of the inflorescence (Corbesier et al. 2007; Walker and Bennett 2023).

*FT*-like genes have expanded by gene duplications occurring independently in nearly all modern angiosperm lineages, including wheat and barley, with 12 FT paralogs (Chardon and Damerval 2005; Faure et al., 2007; Halliwell et al., 2016; Bennett and Dixon, 2021). In barley, *FLOWERING LOCUS T1* (*FT1*), the major floral inducer, is expressed in the leaves, and high transcript levels of *FT1* are linked to early flowering (Yan et al., 2006; Faure et al., 2007; Lv et al., 2014; Digel et al., 2015). *PHOTOPERIOD 1* (*Ppd-H1)* is an upstream regulator of *FT1* orthologous to the *PSEUDO RESPONSE REGULATOR (PRR)* genes of the circadian clock in Arabidopsis (Turner et al., 2005; Beales et al., 2007; Shaw et al. 2013). Two major haplotypes have been described in barley, which differ for a non-synonymous mutation in the conserved CCT (CONSTANS, CO-like, and TOC1) domain (Turner et al., 2005). The ancestral wild-type (WT) allele, predominant in wild barley, landraces, and winter barley, is linked to rapid upregulation of *FT1*, floral development, and flowering under long days (LDs) (Turner et al, Digel et al. 2015). A mutation in the CCT domain of *Ppd-H1* is associated with delayed and reduced upregulation of *FT1* in the leaf and late flowering under LDs and was selected in spring barley, presumably as an adaptation to long growing seasons in northern cultivation areas (Turner et al., 2005; Jones et al., 2008). It has been demonstrated that *Ppd-H1* controls leaf size in barley, which is associated with changes in total cell number and the differential regulation of *FT1* in the leaf (Digel et al. 2016). However, it is not known if *Ppd-H1* controls flowering time, plant architecture and leaf size only through *FT1* or additional genes and networks. Furthermore, the genetic networks downstream of *FT1* in the leaf and SAM and their effects on leaf (source) and inflorescence (sink) development are poorly understood.

In this study, we aimed to dissect the effects of *FT1* on the development of different shoot meristems and organs, and thus on shoot and inflorescence architecture and source-sink relationships during development. We dissected the development of the shoot apical meristem in *ft1* knock-out mutants of the spring barley cultivar Golden Promise, with a mutated *ppd-H1* allele and a derived introgression line carrying a wild-type *Ppd-H1.* Furthermore, we analysed the total transcriptome and carbon metabolites in developing shoot apical meristems and leaves.

*ft1* plants were characterised by substantial changes in the determinacy of different shoot and inflorescence meristems, organ number and size, senescence patterns, and overall plant longevity. The *ft1* mutants exhibited a strong transcriptional reprogramming of genes involved in both the light and dark reactions of photosynthesis, alongside an upregulation of genes associated with sugar catabolism and stress responses. Quantification of soluble sugars and starch in leaves and spikes indicated that the impaired development and growth of *ft1* inflorescences were not due to carbon limitations, but rather to a reduced sink demand, resulting in feedback inhibition of photosynthesis and activation of reactive oxygen species (ROS) scavenging and proteostasis pathways.

## Results

### *FT1* controls the timing of vegetative and reproductive development and plant longevity

Our objective was to investigate how *FT1* affects shoot and inflorescence development in barley, the onset and duration of flowering and transcriptional networks in the leaf and the developing shoot apical meristem. For this purpose, we generated CRISPR-Cas9 induced mutant lines by targeting the coding sequence (CDS) of *FT1* in two different genetic backgrounds: the spring cultivar Golden Promise (GP) carrying the mutated *ppd-H1* allele and GP-fast, which carries a wild-type (WT) *Ppd-H1* allele introgressed from the winter cultivar Igri (Gol et al., 2020; Buchmann et al., 2025). We generated 17 independent M2 lines, 14 in GP-fast and three in GP, as confirmed by sequencing the *FT1* locus and scoring for differences in flowering time. Two late-flowering mutants in the background of GP-fast (*ft1.1, ft1.2*) and one late-flowering mutant in the background of GP (*ft1.3*) were chosen for further experiments. These were characterized by a single base pair insertion at position 114 (+T, *ft1.1*), a deletion at position 113 (-C, *ft1.2*), and a deletion at position 88 (-G, *ft1.3*) (Supplemental Figure S1A). These indels resulted in frameshifts and premature stop codons, reducing protein length from 177 aa (WT) to 59 aa (*ft1.1*), 60 aa (*ft1.2*), and 29 aa (*ft1.3*) (Figure 1A; Supplemental Figure S1B). All three *ft1* mutant alleles lacked the DPDxP and GxHR domains conserved in PEBP proteins (Banfield and Brady, 2000), segment B, and the LYN triad, which form the external loop crucial for the interaction with bZIP transcription factors (Ahn et al., 2006). The truncated *ft1* proteins also lacked the residues Y85 and Q104, important for flowering-activating (Hanzawa et al., 2005; Ahn et al., 2006) as well as the amino acid residues R62, T66, P94, F101, and R130, which are critical for FT-14-3-3 interactions in wheat and rice (Taoka et al., 2011; Li et al., 2015) (Figure 1A). We scored the lines for development and plant architecture traits in two experiments with different pot sizes, small (75 cm^3^ wells) and large (1500 cm^3^), that affected development, architecture and main culm survival.

**Figure 1.**
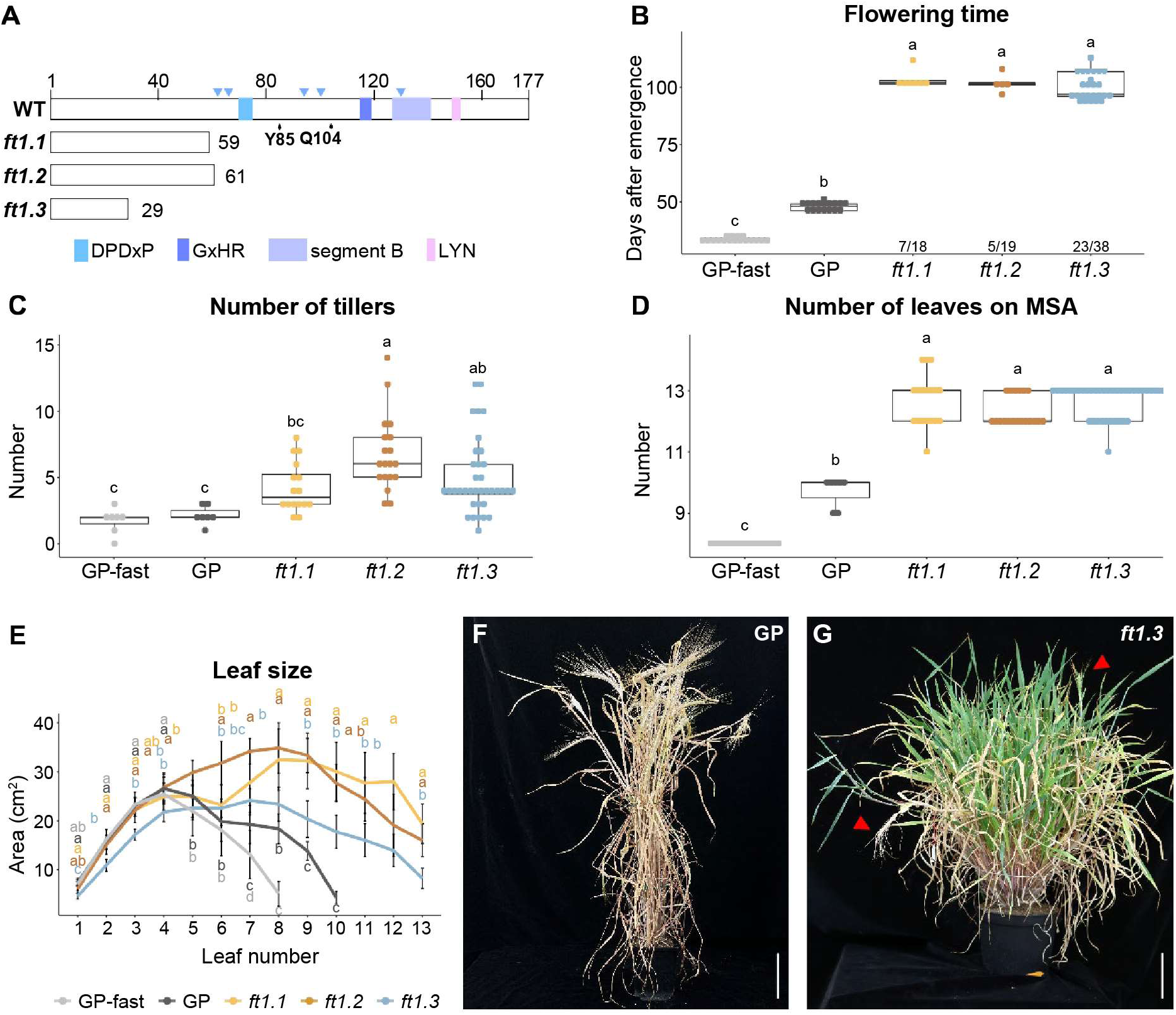
*ft1* mutant alleles and trait variation. **A** Schematic overview of the FT1 protein sequence in *ft1* mutants *ft1.1*, *ft1.2*, and *ft1.3* compared to wild-type (WT). Residues Y85 and Q104, important for flowering-activating function, are marked (Hanzawa et al., 2005; Ahn et al., 2006). The boxes represent DPDxP and GxHR domains highly conserved in PEBP proteins (Banfield and Brady 2000), and segment B and the LYN triad, which form the external loop crucial for interaction with a bZIP transcription factor (Ahn et al., 2006). Blue triangles indicate the amino acid residues R62, T66, P94, F101, and R130, critical for FT-14-3-3 interactions in wheat and rice (Taoka et al. 2011; Li et al. 2015). The numbers on the right indicate the protein length of the mutated FT1 proteins. **B-E** GP-fast, GP, and *ft1* mutants were grown under controlled long-day conditions and different traits related to development and plant architecture were scored at flowering (B, C), the end of stem elongation (D), or during development (E). Flowering was scored for the main culm, or, in case the main culm was aborted, of the first tiller. n = 5-27. Tiller number (C) was scored from 7 WT plants and 16 to 36 mutant plants. **E** Leaf number and leaf size were measured in leaves on the main culm. **F** Leaf size was determined by multiplying leaf length and width. Error bars indicate the standard deviation of the mean. n = 10. Significance levels were determined by one-way ANOVA and subsequent Tukey’s test (*p* ≤ 0.05). F-G Representative images of a mature GP plant (F) at four months after emergence, and a *ft1.3* plant (G) at ten months after emergence with senescing and newly emerging spikes (red arrows) and green leaves. Scales = 10 cm.

The parents GP-fast and GP, and the *ft1* mutant lines *ft1.1, ft1.2,* and *ft1.3* were grown in small pots (75 cm^3^) under controlled long-day (LD) conditions to determine the effect of *FT1* on flowering time and leaf number of the main culm (Zadoks scale; Zadoks et al., 1974), and total tiller number. GP-fast flowered 34 days after emergence (DAE), followed by GP at 48 DAE and the *ft1* mutant lines at 102 DAE (Figure 1B). While GP-fast with a wild-type *Ppd-H1* allele flowered 14 days before GP with a mutant *ppd-H1* allele, we did not observe significant differences in flowering time among the *ft1* mutant lines in GP-fast versus GP (*ft1.1*: 103 DAE; *ft1.2*: 102 DAE; *ft1.3*: 100 DAE) (Figure 1B). These observations suggested that *Ppd-H1* acted only or primarily through *FT1* on flowering time. At the onset of flowering, GP plants had produced, on average, 2.1 secondary tillers, not significantly more than the early-flowering GP-fast plants, which produced, on average, 1.7 tillers (Figure 1C). *ft1* mutants produced significantly more tillers than their respective parent (ft1.1: 4.3; ft1.2: 6.7; ft1.3: 5 tillers) (Figure 1C). We also investigated whether the extended development in the *ft1* mutants affected the number, size and emergence rate of leaves on the main culm. Leaf numbers on the main culm differed significantly between GP-fast, GP and *ft1* mutants (Figure 1D). GP-fast produced an average of eight leaves, GP an average of 10 leaves and the *ft1* mutants 13 leaves on the main culm (Figure 1D). Leaf size increased in the parental lines from leaf one to five and then strongly declined until the flag leaf (Figure 1E). By contrast, in *ft1* mutants, leaf size increased until leaf nine and then declined, so that the sizes of the last leaves were significantly increased in all *ft1* mutants compared to the parents (Figure 1E). These size differences were mainly caused by increased leaf lengths in the mutant lines (Supplemental Figure S2A-B). The *ft1.3* mutant in the background of GP produced leaves that were significantly smaller and thinner than the leaves of the *ft1.1* and *ft1.2* mutants in the background of GP-fast (Figure 1E, Supplemental Figure S2 A-B). The leaf emergence rates were similar for all genotypes during early development, but started to significantly differ from 18 DAE when emergence rates were fastest in GP-fast, followed by GP and then the *ft1* mutants (Supplemental Figure S2C). The number of visible and green tillers of GP-fast and GP increased until the onset of flowering (Supplemental Figure S2D). In contrast, *ft1* mutants continued tillering until 140 DAE when the experiment was stopped (Figure 1F-G). In the *ft1* mutants, a first wave of tillering was followed by a period of tiller abortion and a second wave of tillering after the onset of flowering (Supplemental Figure S2D). The tiller number and emergence rate in *ft1.3* (GP) plants were significantly increased compared to *ft1.1* and *ft1.2* (GP-fast) plants (Supplemental Figure S2D).

In a separate experiment, in larger pots (1.5 L), we tested for effects of *FT1* on plant longevity as well as for effects of transformation and tissue culture on development and plant architecture by including null segregant lines (*ft1.1-null, ft1.2-null, ft1.3-null*), sister plants of the *ft1* mutant lines without a mutation event within the *FT1* sequence. We confirmed the differences in flowering time, leaf number and tillering between *ft1* mutants and parental lines and demonstrated that null segregants did not differ from their respective WT parents (Supplemental Figure S3A-D). However, in the larger pots, all genotypes produced significantly more tillers, resulting in 100% aborted main culms in the *ft1* plants as compared to 25-75% abortion of main shoots in the smaller pots (75 cm^3^) (Supplemental Figure S3E-F). However, GP, GP-fast plants, and null mutants were fully senesced at grain maturity after four months, whereas *ft1* mutant plants continued producing new tillers and flowering until the experiment was stopped after two years (Figure 1F-G). The *ft1* mutant plants did not undergo whole-plant senescence, but senescence was restricted to individual tillers. This resulted in plants with many tillers and a high amount of vegetative biomass (Supplemental Figure S4). In summary, *FT1* had pleiotropic effects on flowering time, tillering, leaf development, and plant longevity. All *ft1* mutants flowered at the same time, suggesting that *Ppd-H1* controls time to flowering primarily through *FT1*. In contrast, leaf size and tiller emergence and number at flowering time differed between *ft1.1 and ft1.2* mutants in GP-fast versus *ft1.3* in the GP backgrounds, suggesting that *Ppd-H1* might control leaf size and tillering through other genes than *FT1.* However, the observed differences might also be controlled by other genetic variation than *Ppd-H1* in the introgression line GP-fast (Buchmann et al. 2025).

### *FT1* accelerates the vegetative and reproductive development and increases fertility

To analyse the effects of *FT1* on the developmental subphases of the main shoot apex (MSA), the MSA of GP-fast, GP, and the three mutant lines *ft1.1*, *ft1.2,* and *ft1.3* were dissected during development and scored according to the scale by Waddington et al. (1983). In addition to the developmental stage, we scored the number of spikelet meristems (SMs) and FMs (floral meristems), and the inflorescence sizes to calculate the rate of SM initiation, FM development, and FM abortion.

The MSA of GP-fast plants developed significantly faster than that of GP; GP-fast plants transitioned to the double ridge stage (W2.0) at 10 DAE compared to 15.5 DAE in GP and 25 DAE in *ft1* mutant plants (Figure 2A). The MSA of GP-fast plants showed a linear development across all Waddington stages. In contrast, GP and *ft1* mutants were further delayed during floral development (Figure 2A). Consequently, GP-fast reached the pollination stage (W10.0) at 38 DAE, GP plants at 52 DAE, and *ft1* plants, on average, at 101 DAE (Figure 2A). No significant differences in the timing of the developmental subphases, the duration of SM induction, and floral development could be observed amongst the *ft1* mutants, regardless of their parental background (Figure 2A). The average rate of SM induction was 0.9 SMs /day in *ft1* mutants and, therefore, strongly reduced compared to the parents with 2.3 SMs /day in GP and 2.9 SMs /day in GP-fast (Supplemental Table S1). While SMs were induced until the same developmental stage (W4.5-W5.0) in both parents and *ft1* mutant lines, they were induced over a longer time period in the *ft1* mutants and GP due to delayed development (Figure 2B). Consequently, the maximum SM numbers on the MSA of *ft1* mutants and GP were significantly higher than GP-fast, even though SM induction rates (SMs/day) were reduced (Figure 2B; Supplemental Table S1). However, the higher number of SMs did not correlate with an increased inflorescence size, as the inflorescence density (measured in SMs/mm inflorescence) was significantly increased in *ft1* mutants and GP compared to GP-fast (Supplemental Figure S5A-C). We then scored floret abortion if SMs or FMs were stalled in development and did not continue to form florets. The total number of SMs/FMs that did not develop into florets was much higher in *ft1* plants compared to parents, with an average of 11.3 aborted SMs or FMs compared to 0.7 in GP-fast and 1.6 in GP on the main spikes (Supplemental Table S1).

**Figure 2.**
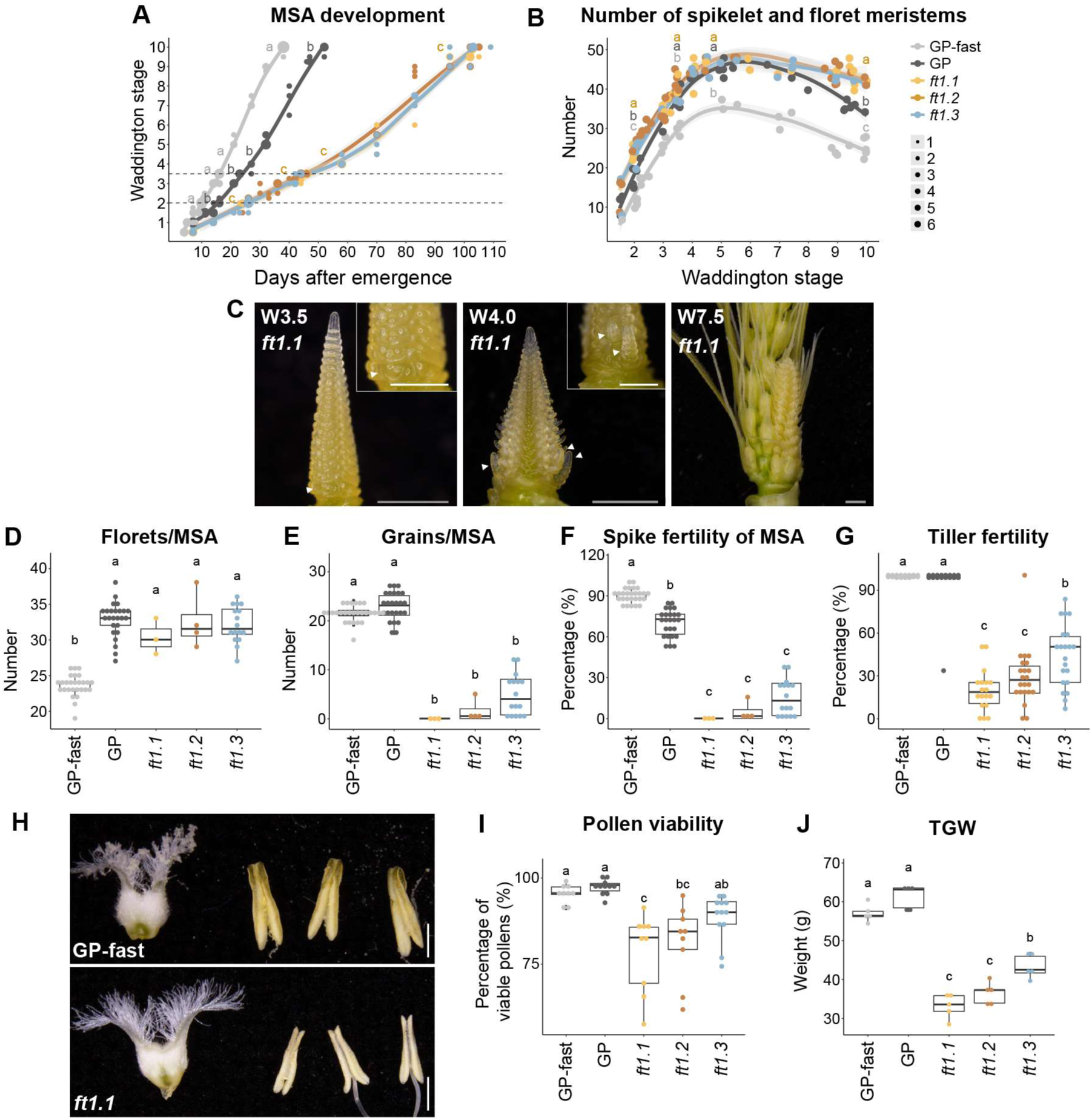
Effects of *FT1* on reproductive development. **A-D** The main shoot apex (MSA) development was monitored from 4 days after emergence to flowering on the main culm of GP-fast, GP, and *ft1* mutants. A Development of the MSA according to the scale by Waddington et al. (1983). Dot sizes indicate the number of plants per data point (1-6), and grey areas show the 95% confidence interval of a polynomial regression (Loess smooth line). The dotted lines indicate the transition from vegetative to reproductive growth (W2.0) and to floral development (W3.5). **B** Number of spikelet and floret meristems on the MSA. Significance levels in (A) and (B) were determined by one-way ANOVA and subsequent Tukey’s test (p ≤ 0.05) at developmental stages W2.0, W3.5, W4.5/W5.0, and W10.0. Values from *ft1* mutants were combined and compared to GP-fast and GP. The colors of the characters indicate the respective group (light grey: GP-fast, dark grey: GP, orange: mutants). **C** Representative MSA of *ft1.1* plants during floral development (W3.5 to W7.5). White arrows indicate secondary inflorescences at the base of the primary inflorescence. A small image in the top right shows the magnified view of the same MSA from a different angle. White scale bars = 500 μm, grey scale bars = 1000 μm. **D-G** Reproductive traits in *ft1* plants. Floret number (D), grain number (E), and floret fertility (F) were scored on the main culm. Floret fertility was calculated by dividing the number of grains by the number of florets. n = 3-27. Tiller fertility was determined by dividing the number of tillers with a spike that holds at least one seed by the final tiller number. n = 18-27. **H** Representative images of GP-fast and *ft1.*1 ovules and anthers; scales = 1000 μm. **I** Pollen viability, with values averaged for each floret. n = 8-23. **J** Thousand-grain weight (TGW) was measured with five biological replicates, each containing pooled grains from three individual plants. n = 5. Significance levels were determined by one-way ANOVA and subsequent Tukey’s test (*p* ≤ 0.05).

The MSAs of GP-fast, GP, and *ft1* mutants were morphologically similar during vegetative and early reproductive developmental stages (W1.0-W3.0) (Supplemental Figure S5D). However, at the stamen primordium stage (W3.5), secondary inflorescences started to emerge from the basal central SMs of the MSA in *ft1* plants, but not in the parental plants (Figure 2C). We observed this in 60–80% of the dissected main culms of *ft1* plants. This usually occurred only on one side of the inflorescences, in rarer cases (about 10%) on both sides. Compared to the MSA, the secondary inflorescences were delayed but appeared to develop normally. They resulted in branched spikes at the base of the spike (Supplemental Figure S6).

Next, we evaluated the effect of *FT1* on the number of spike-bearing tillers and the number of florets and grains on the main spike. The floret number was significantly higher in GP than in GP-fast, while the grain number per main spike was not significantly different between the two genotypes (Figure 2D-E). In *ft1* mutants, the floret number was increased compared to GP-fast but not to GP (Figure 2D). Spike fertility, the number of grains per floret on the main spike, ranged from 0% to 15% in *ft1* mutant plants and was thus significantly lower than in GP-fast (90%) and GP (70%) (Figure 2F). The tiller fertility, the ratio of spike-bearing tillers with at least one grain versus the total number of tillers, was close to 100% in GP-fast and GP (Figure 2G). In contrast, tiller fertility of *ft1.1*, *ft1.2,* and *ft1.3* was significantly lower, averaging only 20%, 29%, and 44%, respectively (Figure 2G). Compared to GP-fast and GP, *ft1* mutant plants produced normal ovules but smaller anthers with fewer and shrivelled non-viable pollen, as demonstrated by pollen viability testing and electron microscope imaging (Figure 2H-I; Supplemental Figure S7). Based on these measurements, pollen viability was reduced significantly from 96–97% in GP-fast and GP to an average of 81%, 78%, and 88% in *ft1.1*, *ft1.2,* and *ft1.3*, respectively (Figure 2I). We thus concluded that the impaired anther and pollen development contributed to the reduction in floret fertility in the *ft1* mutant plants. *FT1* also affected grain morphology as thousand-grain weight (TGW) and two-dimensional grain area were significantly reduced in *ft1* mutants compared to the parents, mainly due to reduced grain width (Figure 2J; Supplemental Figure S8).

In conclusion, *ft1* mutants were characterized by delayed reproductive development, slower but longer SM and FM induction, and increased floret abortion, which resulted in a strong reduction in the number of grains per main spike compared to the parents. This caused a reduced spike fertility, likely affected by a reduced pollen number and viability. Spike architecture in *ft1* plants was altered by the outgrowth of secondary inflorescences from the basal central SMs, resulting in branched spikes. The extended leaf growth, inflorescence meristem activity, and plant longevity observed in the *ft1* mutant lines suggest that *FT1* plays a critical role in establishing the determinacy of various meristem types, thereby coordinating vegetative and reproductive development and ultimately maintaining source–sink homeostasis.

### *FT1* affects the expression of genes involved in stress response, chromatin remodelling and carbon metabolism

To identify genes and genetic networks underlying the *FT1*-controlled phenotypes, we conducted transcriptome profiling in *ft1* mutants, GP, and GP-fast. For this purpose, we harvested leaves and MSAs at key reproductive stages: the initiation of spikelet development (W2.0), the onset of floral development (W3.5), and the end of spikelet induction (W5.0).

Principal Component Analyses (PCA) of the leaf and MSA transcriptome data demonstrated that PC1 separated the genotypes, whereas PC2 separated the developmental stages for leaf and MSA samples, while the genotype and stage-dependent clusters were more pronounced in MSA than leaf samples (Supplemental Figure S9A, S10A). GP with low *FT1* expression levels exhibited transcriptional profiles intermediate between those of the mutants with non-functional *ft1* and GP-fast, characterised by a strong induction of *FT1* expression (Supplemental Figures S10A, S11). We focused our transcriptome analysis on *FT1*-dependent effects, as we found no significant developmental differences in the timing of MSA development and flowering among the *ft1* mutants and wanted to exclude possible effects caused by the background introgression in GP-fast (Figure 1C, 2A). We thus identified differentially expressed genes (DEGs) that were consistently regulated across all three *ft1* mutants relative to their respective parental lines at the same stage. We identified a total of 646 and 628 DEGs in leaf and MSA, respectively (Figure 3A-B; Supplemental Datasets S1-S2). In the leaf, 395 DEGs were upregulated and 251 downregulated, and in the MSA, 396 transcripts were up- and 232 downregulated in the *ft1* mutants (Figure 3A-B). The number of DEGs increased in both tissues with the developmental stage (Figure 3A-B). Only 40 DEGs overlapped between leaf and MSA tissues, which included the developmental genes *ODDSOC1* (*HvMADS25*), *AGL14,* and the starch-degrading enzyme *BETA-AMYLASE*, which were strongly upregulated in both leaf and MSA in *ft1* mutants (Supplemental Figures S11, S12).

**Figure 3.**
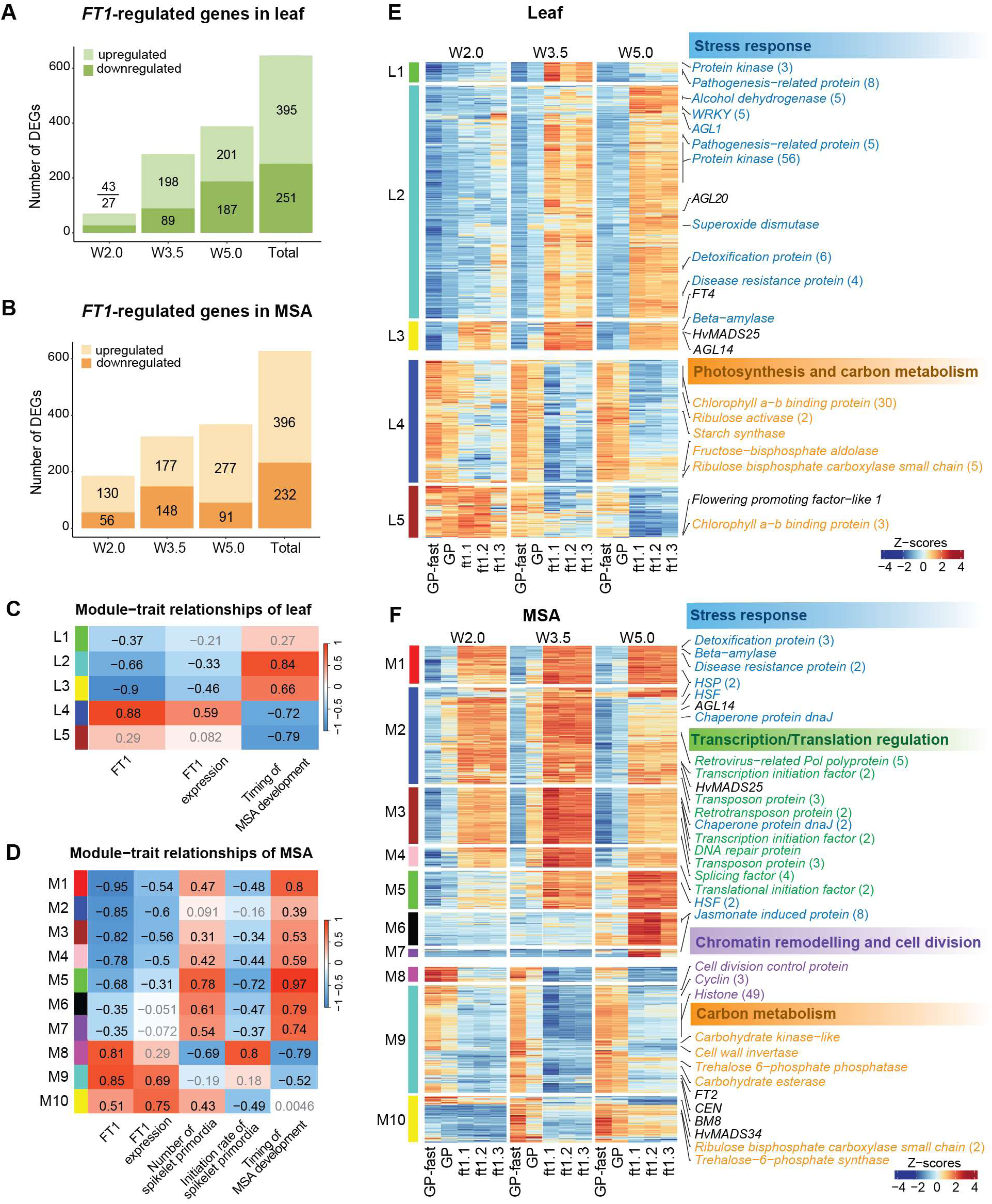
Effects of *FT1* on transcriptomic changes in leaf and inflorescence. **A-B** The number of differentially-expressed genes (DEGs) regulated by *FT1* at each developmental stage and in total across all stages in the leaf (A) and main shoot apex (MSA). Genes that were either up- or down-regulated in all *ft1* mutants (*ft1.1 vs.* GP-fast, *ft1.2 vs.* GP-fast, and *ft1.3 vs.* GP) at the same stage were considered as *FT1*-regulated genes. **C-D** Correlation between *FT1*-regulated genes and genetic and phenotypic traits. L1-5 are modules identified in leaf (C), and M1-10 are modules in MSA (D). Values in the cell represent the correlation coefficient. Grey values show non-significant correlations. The color scale on the right side indicates the strength and direction of the correlation. Details can be found in Supplemental Figure S13. **E-F** The heatmaps show the transcript profiles of genes in each module. Top gene ontology (GO) terms for modules are shown on the side of the heatmap with representative genes in the GO terms. Genes involved in stress response (blue), photosynthesis and carbon metabolism (orange), transcription and translation regulation (green), and chromatin remodelling (purple) are labelled next to their transcript heatmaps. Details can be found in Supplemental Datasets S1 and S2. The color scale represents the range of Z-score normalized mean transcript per million (TPM) of three biological replicates.

To identify co-expression clusters correlated to genetic and expression variation of FT1, the timing of MSA development and SP and FM induction, a weighted gene co-expression network analysis (WGCNA) was performed in the leaf and MSA, respectively (Figures 3C-D; Supplemental Figure S13). We identified five co-expression clusters in the leaf and ten in the MSA. Co-expression clusters with transcripts upregulated in the *ft1* mutants were enriched for functions in stress response in both leaf and MSA, and for functions in transcription and translation only in the MSA (Figures 3E-F; Supplemental Figures S9B, S10B). Co-expression clusters downregulated in the *ft1* mutants were enriched for functions in photosynthesis and carbon metabolism in the leaf and MSA, and for chromatin remodelling only in the MSA (Figures 3E-F; Supplemental Figures S9B, S10B; Supplemental Datasets S3, S4).

In the leaf, co-expression module L2 (325 DEGs), with the strongest positive correlation to the timing of MSA development and negative correlation to FT1, included 79 genes with putative functions in the mitigation of oxidative stress (reactive oxygen species (ROS) scavengers, *CATALASE*), heavy metal toxicity (*HEAVY METAL TRANSPORT/DETOXIFICATION SUPERFAMILY PROTEINS* [*DTX*], *ABC TRANSPORTER*s), heat (*HEAT SHOCK PROTEIN*s [*HSP*s]) and pathogens (*HYPERSENSITIVE-INDUCED RESPONSE PROTEIN* [*HIR*]), *NBS-LRRs*) (Figures 3C, 3E; Supplemental Figure S11; Supplemental Dataset S1). In addition, 143 genes were associated with metabolic adjustment and stress-related signalling pathways. These included kinases (*SERINE/THREONINE* [*Ser/Thr*] kinases, *RECEPTOR-LIKE KINASES* [*RLKs*]), *PHOSPHATASE* 2C genes (*PP2C*s), transcription factors (*WRKY*, *NAC*, *MYB*, *bZIP*, and *bHLH* families), transporters (*SUGAR TRANSPORTER* genes, *PHOSPHATE TRANSPORTER 1*, and *NRT/PTR* family transporters), as well as enzymes (*FRUCTOKINASE FRK2 KINASE*, *BETA-AMYLASE*, *GLYCOSYLTRANSFERASE,* and *BETA-GLUCOSIDASE*) involved in carbon catabolism under stress conditions (Figure 3E, Supplemental Figure S11; Supplemental Dataset S1). In addition, the floral repressor *FT4* and developmental regulator AGL20 were strongly upregulated in the *ft1* mutant plants but were not or lowly expressed in GP and GP-fast (Figure 3E; Supplemental Figure S11). In contrast, leaf module L4 (171 DEGs), which showed the strongest negative correlation with the timing of MSA development, was enriched for genes involved in photosynthesis and carbon fixation (Figure 3E; Supplemental Figures S9B). Specifically, *CHLOLORPHYII A-B BINDING* (*CAB*) genes, *PHOTOSYSTEM I SUBUNIT* genes (*PsaO*, *PsaE-2*), *PHOTOSYSTEM II SUBUNIT* genes (*PsbR*, *PsbW*), *RIBULOSE BISPHOSPHATE CARBOXYLASE SMALL CHAIN 1A* (*RBCS1A*) genes, *RUBISCO ACTIVASE* (*RCA*), and *D-RIBULOSE-5-PHOSPHATE-3-EPIMERASE* (*RPE*) gene were downregulated in the *ft1* mutants (Supplemental Figure S11; Supplemental Dataset S1). These results showed that co-expression clusters in the leaf associated with FT1 function and MSA development were enriched for functions in stress responses, photosynthesis and carbon metabolism.

In the MSA, modules M1–M7 comprising 396 DEGs upregulated in the *ft1* mutant, were, like the leaf modules L1-L3, enriched for stress response genes, such as heat stress responses (*HEAT SHOCK FACTORs* [*HSF*s], *HSP*s), ROS scavenging (*GLUTATHIONE TRANSFERASEs* [*GSTs*], *GLUTAREDOXIN*, *LIPOXYGENASE*s), hormonal stress signaling (*ETHYLEN-RESPONSIVE TRANSCRIPTION FACTOR*s [*ERF*s], *JASMONATE-RESPONSIVE PROTEIN*s, *AUXIN RESPONSE FACTOR 1* [*ARF1*], *PP2C*), detoxification (*DTX*s), ubiquitin-mediated protein turnover (*F-BOX* proteins, *POLYUBIQUITIN*), metabolic stress or starvation responses (*PYRUVATE DECARBOXYLASE*, *PHOSPHOGLYCERATE KINASE*, fatty acid oxidation enzymes, *HEXOSYLTRANSFERASE*s), and programmed cell death (*DEVELOPMENTAL AND CELL DEATH* gene [*DCD*] (Figure 3F; Supplemental Figure S12; Supplemental Dataset S2). Additionally, DEGs in M1–M7 were enriched for functions in transcriptional and post-transcriptional regulation (*TRANSCRIPTION ELONGATION FACTOR TFIIS*, *TRANSLATION ELONGATION FACTOR 1-α*, *SPLICING FACTOR*s), and for transposable elements (CACTA, *RETROVIRUS-RELATED* POL *POLYPROTEIN*s), indicating elevated transcriptional activity in *ft1* MSAs (Figure 3F; Supplemental Figure S12; Supplemental Dataset S2). Moreover, the expression of *MADS-box* gene *ODDSOC2,* flowering repressor *CONSTANT 9-LIKE*, and circadian clock gene *PRR73/ PRR95*, were strongly upregulated in the MSA of *ft1* mutants (Supplemental Figure S12; Supplemental Dataset S2). Notably, modules M1–M7 were negatively correlated with SP numbers and rate of SP initiation, suggesting that delayed SP and FM development was linked to molecular stress response and transcriptional reprogramming in the MSA (Figures 2A, 3D). By contrast, three modules (M8–M10) were negatively correlated with the genetic and expression variation of *FT1* in the leaf. These modules included the *FT1* paralog *FT2*, *MADS-box* transcription factors *CENTRORADIALIS* (*CEN*) and *BM8*, which promote floral development and floret fertility; and genes associated with chromatin remodeling and cell division, such as *HISTONE* and *CYCLIN* family members (Figure 3F; Supplemental Figure S12; Supplemental Dataset S2). Moreover, modules M8–M10 were enriched for genes involved in carbon metabolism and signaling, including *TREHALOSE-6-PHOSPHATE SYNTHASE 1* (*TPS1*) and *TREHALOSE-6-PHOSPHATE PHOSPHATASE G* (*TPPG*), two enzymes that act sequentially to regulate the trehalose-6-phosphate (T6P) balance, as well as *CELL WALL INVERTASE 2* (*CWINV2*), which controls sucrose cleavage in sink tissues (Supplemental Figure S12).

Taken together, the delay in MSA development and decrease in SP and FM induction rates in the *ft1* mutants were linked to a transcriptional stress response and downregulation of transcripts involved in floral development and in carbon assimilation and signalling in leaves and MSA.

### *FT1* affects soluble sugar and starch levels in the leaf and spike

We then tested if the *FT1*-dependent regulation of photosynthesis-related genes in the leaf and of carbon signaling and metabolism in the MSA was linked to changed carbon availability. Therefore, we measured starch and soluble sugars at the end of the day (EOD) and end of the night (EON) in leaves and MSAs at the pre-anthesis stage (W8.0-W8.5). In the leaf, sucrose was most abundant, followed by starch, glucose and fructose at the end of the day, while starch levels were relatively higher than sucrose at EON (Figure 4A). In the spike, sucrose was more abundant than starch, followed by glucose and fructose at EOD and EON (Figure 4A). In the leaf, GP-fast accumulated higher levels of sucrose, glucose and fructose, particularly at EOD, compared to GP and the *ft1* mutants, while GP and *ft1* mutants did not differ for these metabolites in the leaf (Figure 4B). Leaf starch levels were consistently different between GP-fast versus GP and *ft1* mutants at EOD or EON (Figure 4B; Supplemental Figure S14A). In the spike, GP and the *ft1* mutants had higher levels of sucrose and starch at EOD and EON compared to GP-fast (Figure 4B). Furthermore, all three *ft1* mutants displayed higher fructose, glucose and total sugar levels at EON than EOD, while GP-fast and GP sugar levels were not different or lower at EON than EOD in the spike (Figure 4B; Supplemental Figure S14B).

**Figure 4.**
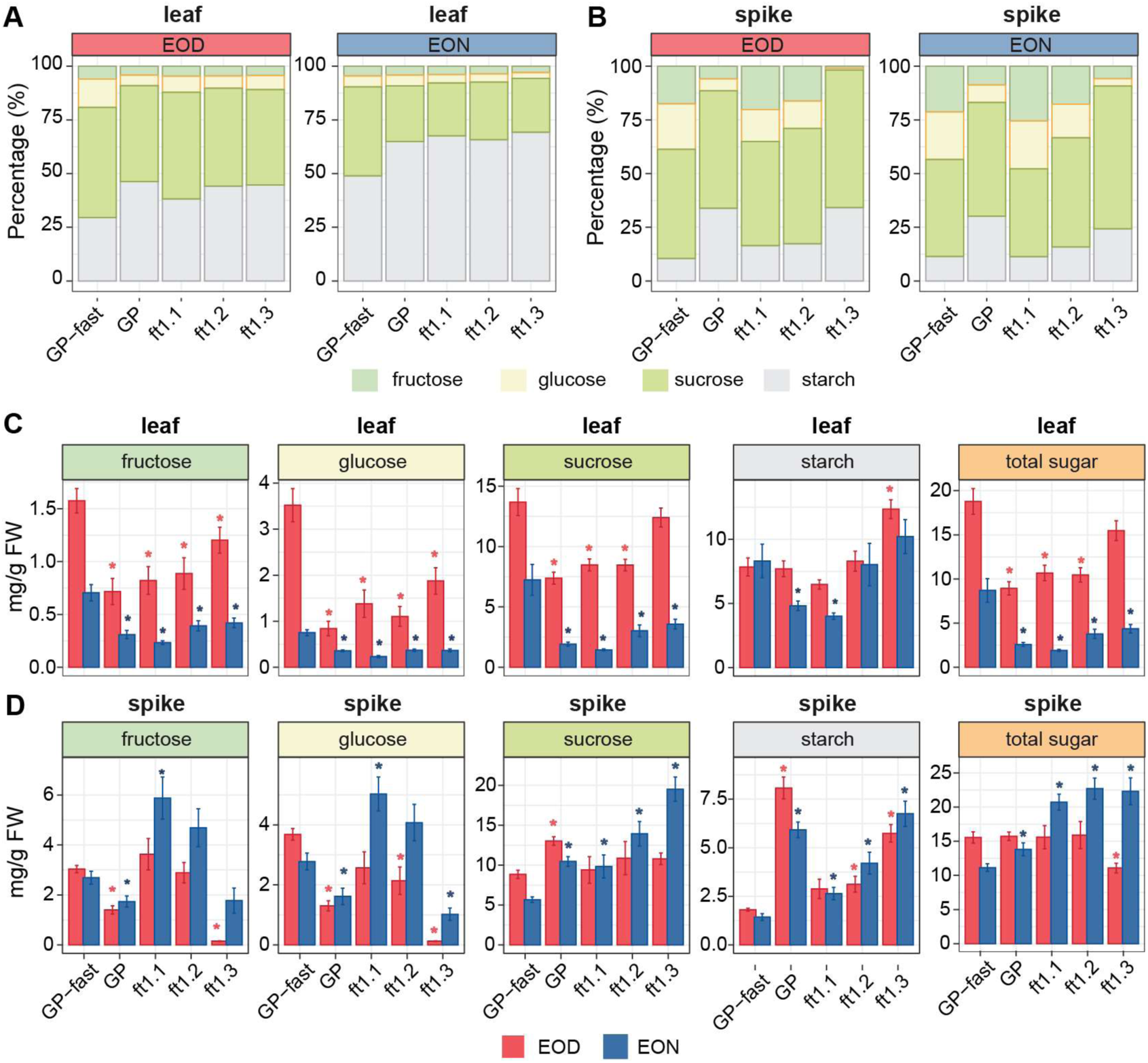
Effect of *FT1* on soluble sugars and starch content in the leaf and spike. **A-B** Proportion of soluble sugars and starch in the leaf (A) at 16 days after emergence and in the spike (B) at developmental stage W8.0-8.5 (before pollination) of GP-fast, GP, *ft1.1*, *ft1.2* and *ft1.3* at the end of day (ZT 16; EOD) and the end of night (ZT 0; EON). **C-D** Soluble sugars and starch content in leaves (C) and spikes (D) at EOD (red) and EON (blue). FW, fresh weight. Error bars indicate the standard error of ten biological replicates. Significance levels were determined by comparing GP or *ft1* mutants to GP-fast at either EOD or EON using Fisher’s F-test for homoscedasticity, followed by Welch t-statistic. Red asterisks indicate significant differences at EOD, blue asterisks at EON. *, *p* ≤ 0.05.

Taken together, *ft1* mutant plants showed reduced levels of soluble sugars in the leaf, but increased sucrose and starch in the inflorescence compared to GP-fast, as well as higher fructose, glucose and total sugar levels in the spike at EON than EOD.

## Discussion

### FT1 coordinates source-sink growth

Loss of a functional FT1 protein not only delayed the transition from vegetative to reproductive development but also slowed down and extended the period of reproductive growth (Figure 1G, 2A). In all three *ft1* mutant lines, the transition to reproductive development and the timing of reproductive phases were consistently delayed, irrespective of the allelic state of *Ppd-H1*, indicating that *Ppd-H1* regulates MSA development primarily via FT1 (Figure 2). However, leaf size and tiller number at flowering time differed between *ft1.1* and *ft1.2* mutants in the GP-fast background versus *ft1.3* in the GP background, suggesting that *Ppd-H1* or other variants introgressed from Igri into GP-fast might control these traits (Figures 1, 2, 4; Supplemental Figure S15).

In all *ft1* mutants, prolonged vegetative development led to the formation of more leaf primordia, increased leaf numbers, and enhanced tillering, thereby increasing source strength (Figure 1). Similarly, the prolonged activity of the inflorescence meristem resulted in an extended period of spikelet initiation and an increased number of spikelets on the main inflorescence (Figure 2). In addition, basal central SMs reverted into secondary inflorescence meristems, giving rise to a branched spike architecture (Figure 2; Supplemental Figure S6). The reversion of spikelet meristems to secondary inflorescences was also observed in the barley *mnd1* mutant and wheat and barley lines, with high expression levels of *SHORT VEGETATIVE PHASE 1* (*SVP1*)-like genes (Trevaskis et al. 2007; Walla et al. 2020; Li et al. 2021). Similarly, the upregulation of the *SVP-like* gene *ODDSOC1* (*HvMADS25*) in the leaf and the inflorescence was linked to floral reversion in the *ft1* mutants (Figure 2; Supplemental Figures S11, S12). Furthermore, the downregulation of floral homeotic genes such as *FT2* and *BM8*, which are associated with floral organ development in wheat and barley (Digel et al. 2015, Shaw et al. 2019, Mulki et al. 2018), correlated with the arrested floral development and pronounced reduction in grain number per spike in the *ft1* mutant lines (Figure 2; Supplemental Figure S12). Consequently, sink strength was strongly reduced in *ft1* mutant plants despite an increase in spikelet primordia numbers during early reproductive growth.

The *ft1* mutants exhibited prolonged leaf development. Previous work identified *Ppd-H1* as a major determinant of leaf size in barley (Digel et al., 2016). Our findings suggest that *Ppd-H1* influences leaf size by regulating *FT1* expression in leaves. *FT1* may affect leaf size either directly, by modulating leaf determinacy and cell proliferation, or indirectly, by altering the timing of inflorescence development and thus stem elongation (Figure 2). The latter is closely linked to the phyllochron, the interval between the emergence of successive leaves, which determines how long a leaf can grow before the next leaf emerges (Digel et al. 2016) (Supplemental Figure S2). Additionally, *ft1* plants exhibited increased tillering. Similar effects of *FT* homolog knockouts on branching have been reported in other plant species. In wheat, loss-of-function or deletion of the *FT-B1* allele resulted in increased leaf and tiller numbers (Dixon et al., 2018; Finnegan et al., 2018). In tomato, mutations in *SFT*, the *FT* ortholog, enhance leaf production and branching (Park et al., 2014). Likewise, *Cas9-GhFT* plants in cotton exhibit indeterminate meristem growth and altered branching architecture (Sang et al., 2025).

The *ft1* mutants exhibited prolonged tillering over many months and lacked whole-plant senescence, suggesting that *FT1* controls not only the onset, but also the duration of flowering and plant longevity (Figure 1; Supplemental Figures S3, S4). This growth habit is reminiscent of perennial plants, which do not undergo whole-plant senescence but maintain undifferentiated meristems (Albani and Coupland, 2010). Similarly, Melzer et al. (2008) showed that the downregulation of flowering time inducers in annual Arabidopsis resulted in plants with recurrent growth cycles and markedly increased longevity. Recent studies in *Arabidopsis thaliana* have shown that flowering time genes regulate not only the initiation but also the duration and termination of flowering (Balanzà et al., 2018; Miryeganeh, 2021; González-Suárez et al., 2023). Consistent with this, our findings suggest that *FT1* controls both the onset and duration of flowering, as well as senescence patterns, thereby influencing overall plant longevity in barley.

### Reduced sink growth is linked to a feedback inhibition of photosynthesis genes, and activation of stress response pathways

The changes in source-sink tissues observed in *ft1* mutant lines were accompanied by extensive transcriptional reprogramming, the downregulation of genes involved in both the light and dark reactions of photosynthesis, as well as the upregulation of genes associated with sugar catabolism and stress responses (Figure 3; Supplemental Datasets S1, S2). Specifically, transcripts encoding photosystem components and RuBisCO subunits were reduced, while genes involved in starch breakdown and the utilisation of cell wall-derived sugars were upregulated (Supplemental Figures S11, S12). Accordingly, soluble sugars, particularly sucrose, were reduced in the *ft1* mutant leaves. In contrast, sucrose and starch levels were rather increased in *ft1* mutant spikes compared to GP-fast. Specifically, the increase of fructose and glucose at EON compared to EOD in the MSA likely reflected reduced carbon use caused by impaired spike development and growth, as indicated by the downregulation of cell cycle genes in *ft1* inflorescences (Figure 4). Furthermore, low *TPS1* expression likely reflected reduced T6P signalling, thereby promoting sugar breakdown as seen in the strong upregulation of *BETA-AMYLASE* expression and suppressing growth processes (Fichtner and Lunn, 2021; Kumar et al., 2023) (Supplemental Figures S11). These results suggest that the delayed growth and development of *ft1* inflorescences were not caused by carbon limitations, but rather by changes in development and growth, and thus sink strength.

Among the most striking molecular alterations observed in *ft1* mutants was the pronounced upregulation of stress-response genes in both leaf and MSAs tissues (Figure 3; Supplemental Datasets S1, S2). The upregulation of *CHAPERONEs* and *HSPs*, together with genes involved in reactive oxygen scavenging, and in ubiquitin-mediated protein turnover, was a strong indicator of the unfolded protein response (UPR), reflecting cellular stress due to misfolded proteins and activation of proteostasis pathways (Sewelam et al., 2016) (Figure 3; Supplemental Datasets S1, S2). Concurrently, the downregulation of *HISTONE* genes and upregulation of transcripts related to transcription and translation suggested active chromatin remodelling, likely facilitating enhanced access to and transcription of stress-responsive genes in the inflorescence (Figure 3; Supplemental Dataset S2). The observed transcriptional stress signature mirrored the molecular responses reported in Arabidopsis and barley shoot apical meristems under high-temperature conditions, where strong induction of HSFs and HSPs was observed (Ejaz et al. 2017; Lan et al., 2025; John et al., 2022, 2024; Olas et al., 2021). In rice, the photoperiod regulator *HEADING DATE 1 (Hd1)*, an upstream activator of FT under long-day (LD) conditions, has been implicated in modulating salt stress sensitivity, suggesting that photoperiodic flowering regulators may play broader roles in abiotic stress adaptation (Biancucci et al. 2025). On the other hand, it has been demonstrated that low sink demand can lead to the downregulation of the photosynthetic machinery and the generation of ROS, which in turn triggers the induction of protective mechanisms, such as antioxidant enzymes, to prevent damage (Paul and Foyer, 2001, Lv et al., 2020). We thus speculate that an increase in photosynthetically active source tissue and a reduction in sink tissue in the *ft1* mutant plants might have caused a metabolic feedback inhibition of photosynthesis genes and activation of stress signalling pathways.

Our work reveals that *FT1* coordinates the development of vegetative and reproductive meristems and organs with plant physiology and metabolism, thereby regulating source–sink relationships and balancing plant longevity with reproductive output. Identification and characterisation of metabolites and signals controlling the *FT1*-dependent source-sink changes will be important for manipulating the relationship between plant longevity and reproductive output.

## Acknowledgements

We greatly thank Rebekka Schüller, Nina Döring and Sabine Sommerfeld for excellent technical assistance, and Dominik Brilhaus for data management support.

## Contributions

GH and MvK conceived and designed the experiments. GH genotyped and identified the *ft1* mutants, conducted plant phenotyping, sampled for RNA-seq, and analysed and interpreted the data. TL assisted with sample collection and analysed the RNAseq data. EHB contributed to RNAseq analysis. TR extracted and analysed carbohydrates with the help of PW and KW. JK and GöH designed and cloned Cas9 transformation vectors and performed the transformation and regeneration of *ft1* mutants. RS provided guidance, feedback, and support throughout the project. GH and TL wrote the manuscript with the help of MvK.

## Materials and Methods

### Plant material

Spring cultivar Golden Promise (GP, carrying a mutated *ppd-H1* allele) and its derived introgression line GP-fast (wild-type *Ppd-H1* introgressed into GP from Igri (Gol et al., 2021; Supplemental Figure S15) were transformed using a method previously described (Marthe et al. 2015) with an FT1-specific guide (CRISPR-g)RNA/Cas9 construct (pGH465, Supplemental Figure S16) that directly derives from the generic binary vector pSH121 (Gerasimova et al. 2016) to create plants with a nonfunctional *FT1 (HORVU.MOREX.r3.7HG0653910)* gene. The sgRNA targeted a sequence within the first exon of *FT1* (position 97-116 relative to the start codon, 5’→3 ‘: TGACCTTCGGGAACAGGGCCGTGTCCAA). M1 grains were grown in the greenhouse for single-seed propagation.

From 17 different M1 lines, 2-3 grains were grown in a plant growth chamber under controlled LD conditions as described below. DNA was extracted from leaf material using the KingFisher Flex (ThermoFisher) and the BioSprint 96 DNA Plant Kit (QIAGEN) according to the manufacturer’s instructions. To identify mutations within the CDS of *FT1*, the complete genomic sequence of *FT1* was amplified with flanking primers (fwd 5’→3’: GAAGGAAGGAGAAATGGCCG, rev 5’→3’: GATCGAGCGAGCATTAGTCA). PCR products were cleaned using the ExoSAP-IT PCR Product Cleanup Kit (ThermoFisher) and sequenced with Sanger Sequencing. Sequences were compared using MEGA-11 (Version 11.0.10, Tamura et al., 2021).

Three T-DNA-free lines with homozygous single nucleotide polymorphisms (SNPs) were chosen for further experiments and termed *ft1.1*, *ft1.2* (in the background of GP fast), and *ft1.3* (in the background of GP). T-DNA-free sister plants derived from the three mutant lines and that did not show a mutation event within the FT1 sequence were chosen as null segregant lines.

### Growth conditions and plant phenotyping

All plants were grown in soil in controlled growth chambers under long days (LD, 16 h light, 20 °C, PAR ∼250 µmol/m^2^s; 8 h dark, 16 °C). Plants were grown in Einheitserde ED73 (Einheitserde Werkverband e.V.) with 7% sand and 4 g/L Osmocote Exact Hi.End 3-4M, 4^th^ generation (ICL Group Ltd.). All plants were stratified for 3–4 days at 4 °C and darkness after sowing.

GP-fast, GP, *ft1.1*, *ft1.2,* and *ft1.3* plants were grown in QuickPot 96T trays (HerkuPlast Kubern GmbH, pot volume 75 cm^3^). At least ten plants were scored per genotype. Flowering was scored as the days between the emergence of the seedling from the soil and plants reaching Zadoks’ stage 49 when the awns exited the leaf sheath (Zadoks et al., 1974). Plant height was measured at flowering as the distance between the soil and the flag leaf ligule of the main culm. Tiller number was counted as all secondary tillers that had emerged after the main culm at flowering, and the leaf number was determined on the main culm. Leaf width and length were measured on fully elongated leaves on the main culm as the leaf blade length (from the ligule to the leaf tip) and the maximum width of the blade. The emergence of leaves and tillers was scored every 2-3 days by visual inspection of the plants. As soon as tillers were visibly aborted (turned yellow), they were excluded from the scoring. If available, floret and grain number were determined on the main culm. This was used to calculate the spike fertility (as grains per florets). Tiller fertility was determined by dividing the number of tillers with a spike that held at least one grain by the total tiller number. The main culm abortion rate was determined by scoring whether the main culm produced a spike or not.

To test the effect of tiller number on main culm fertility, WT plants, mutant lines and null segregant lines were sown in QuickPot 96T trays (HerkuPlast Kubern GmbH, pot volume 75 cm^3^). At ten days after emergence (DAE), at least four (null segregant lines) or eight (*ft1* lines) plants per genotype were repotted to single 1.5 L pots, and plants were cultivated under LD conditions for phenotyping. Flowering, plant height, tiller number, and leaf number were scored as described above and the main culm abortion rate was determined for all genotypes

Main shoot apex (MSA) development of GP, GP-fast, *ft1.1, ft1.2,* and *ft1.3* was monitored under LD. Every 4–13 days, the main culms of 3–4 individual plants were dissected, starting 4 DAE for GP-fast and 7 DAE for the other genotypes. Parent plants were dissected more frequently than *ft1* mutants due to the faster development. The stage of the MSA was documented using the stereo microscope Nikon SMZ18 with a Nikon DS-Fi2 camera, analyzed with the NIS-Elements Software (version 5.21.03, Nikon Instruments Europe BV), and quantified according to the Waddington scale (Waddington et al., 1983). This scale rates the progression of spikelet meristem (SM) initiation and the development of the most advanced floret meristem (FM) and pistil of the main inflorescence. During early development (W1.0), the MSA is vegetative, and leaf primordia are initiated. The MSA transitions to a reproductive inflorescence at the double ridge stage (W1.5 - W2.0), when SMs become visible adjacent to the leaf primordia, forming the characteristic “double ridges”. Leaf primordia are suppressed, and instead, SMs are induced until approximately W5.0, which determines the maximum number of spikelets, florets and grains per spike (Digel et al., 2015; Zhong et al. 2021; Thirulogachandar and Schnurbusch, 2021). Floral organ primordia start to differentiate at the stamen primordium stage (W3.5) when the central SM has differentiated into three stamen primordia. At W5.0, the last floral organ, the ovule, emerges. Floral organs grow and develop into florets until anthesis and pollination at W10.0.

Inflorescence size was scored as the distance between the node of the lowest SM and the tip of the inflorescence meristem (IM). The number of developing SMs, including those that had initiated FMs or developed into florets, was determined from Waddington stage W2.0 to W10.0. This data was used to calculate the inflorescence density (SMs per mm inflorescence length). The maximum SM stage (Waddington stage) was determined by plotting the number of SMs against developmental stage (Waddington Stage) or days after emergence (, DAE) and calculated with the R package segmented as the break-point of two separate linear regressions. SM initiation rate equals the slope of the first regression, and the floret meristem (FM) abortion rate is the slope of the second regression. The maximum SM and final FM number were calculated with the linear models provided by segmented. Aborted FMs were calculated by subtracting the final FM number from the maximum SM number. All numbers were rounded to one decimal place.

### Pollen, anther, and grain measurements

Spikes with central florets at approximately Waddington stage W10.0 were used to test pollen viability according to a modified protocol from Peterson et al. (2010). Main culm spikes were harvested from at least three individual plants per genotype. Six central florets (from both sides of the two-rowed spike) were opened from each spike, and anther size was determined by measuring the length of the three anthers per floret using the software Fiji (Schindelin et al., 2012). From 2–3 of these florets, the anthers were transferred into the staining solution (as described in Peterson et al., 2010) without prior fixing. Samples were incubated for 40 min at 100 °C. Then, free pollen in the staining solution were transferred to a microscope slide. The examination was performed using a Nikon stereo microscope (Nikon SMZ18), and the images were taken with a Nikon DS-Fi2 digital camera connected to the microscope. Pollen viability was determined by visually inspecting whether pollen were stained purple (classified as viable) or stained light blue (classified as non-viable). For each floret, 60-400 pollen grains were classified. The viability (ratio of viable to all pollen) was averaged per floret. To determine the average pollen diameter, ten randomly chosen fertile pollen were measured per floret, using Fiji. Numbers were averaged for each floret.

For SEM imaging, pollen were collected from anthers of GP-fast and *ft1.1* plants and transferred into a 2 ml tube with 1 ml of 1x PBS (pH 7.4). The PBS was discarded, and 1 ml fixative solution (1% Glutaraldehyde, 4% Paraformaldehyde, 0.03% Triton-X100, 1x PBS pH 7.4) was added under the fume hood. A vacuum was applied for 1–2 h at room temperature (RT) until the pollen had sunk to the bottom of the tube. The fixative was discarded and followed by three 15-minute washing steps with 1 ml of 1x PBS. The pollen were dehydrated by incubating them step-by-step in an increasing percentage of ethanol for 30 min each: Pollen were transferred to 10% EtOH first, followed by 20%, 30%, 50%, 70%, 90%, and final 100%. Next, pollen were transferred to a 1:2 hexamethyldisilazane (HMDS) solution in 100% EtOH and incubated for 20 min at RT. Subsequently, pollen were transferred to a 2:1 solution of HDMS in 100% EtOH and incubated for 20 min. Finally, pollen were transferred to 100% HMDS and left overnight in the fume hood to dry. Until imaging, pollen were stored dry. SEM images were taken at the Forschungszentrum Jülich.

Different yield parameters were measured using a MARViN ProLine (MARViTECH GmbH) and an external scale on grains from GP, GP-fast, *ft1.1, ft1.2,* and *ft1.3*. Per genotype, five replicates, each containing two grains from three individual plants (thus, six grains in total), were measured with the palea facing upwards. Thousand-grain weight (TGW), grain area, length, and width were determined. Two randomly selected grains per genotype were photographed with the palea and with the lemma facing upwards.

### RNA sample preparation and RNA Sequencing

GP, GP-fast, *ft1.1*, *ft1.2,* and *ft1.3* plants were sown in QuickPot 96T trays (HerkuPlast Kubern GmbH, pot volume 75 cm^3^) and transferred to a plant growth chamber after stratification and cultivated under LD conditions as described above.

For RNA Sequencing, plants were sampled at three developmental stages (W2.0, W3.5, and W5.0). Samples were taken at Zeitgeber Time (ZT) 14–15, shortly before the onset of the night when *FT1* and *Ppd-H1* expression was high. Three replicates were obtained for each developmental stage and each tissue. MSAs were collected under a stereo microscope to ensure the correct developmental stage, and the MSAs of multiple plants (W2.0: 15 plants, W3.5: 5 plants, W5.0: 4 plants) were pooled for one replicate. Leaves were sampled at the same developmental stages, and the material of two different plants was pooled for one replicate. The middle section of the youngest, fully elongated leaf was sampled, resulting in some variation in leaf number across the genotypes due to the differences in development (see Supplemental Table S2). All samples were immediately frozen in liquid nitrogen and stored at -80 °C.

RNA extraction was performed with the RNeasy Plant Mini Kit (QIAGEN) according to the manufacturer’s instructions. Remaining DNA was removed using the RNase-Free DNase Set (QIAGEN). The quantity and quality of the RNA was determined with a Nanophotometer (Implen) and on a 1% agarose gel. Paired-end sequencing was performed by Novogene Co, Ltd. using a NovaSeq PE150 platform (Illumina), resulting in 33–69 million reads (5– 10.5 Gbp) per sample.

### RNA Sequencing analysis

The initial quality control of the raw reads was performed with FastQC and then summarized with MultiQC (version 1.7, Ewels et al., 2016). No trimming of the reads was required. Reads were mapped against the most recent reference transcriptome BaRTv2 (Coulter et al., 2022). Mapping was performed using Salmon (version 1.9.0, Patro et al., 2017), and the mapping rate averaged 90.3% across samples. Genes with at least five counts per million in at least three samples across all genotypes were considered as expressed. Out of 39434 annotated genes in the BaRT2 transcriptome reference, we identified 17459 (44%) and 16930 (43%) genes expressed in MSA and leaf, respectively. The 3D RNA-seq pipeline was used to calculate Transcripts per million (TPM) to generate PCAs (Guo et al., 2021). The false discovery rate (FDR, BH adjusted) was calculated using the R package *edgeR* (Robinson et al., 2010). The log2 fold change (log2FC) was calculated with a pseudo count of 1 and by pairwise comparison of each *ft1* mutant to their respective parent (*ft1.1* and *ft1.2* against GP-fast, *ft1.3* against GP). This was done individually for each tissue and developmental stage. The raw data, including all FDR values, can be found in Supplemental Datasets S1 and S2.

Differentially expressed genes (DEGs) were defined for each tissue as those genes that showed significant differences in expression (FDR < 0.01) in all pairwise comparisons (*ft1.1* vs. GP-fast*, ft1.2* vs. GP-fast and *ft1.3* vs. GP). Since developmental differences were quantitative among GP-fast, GP with low *FT1* expression and the *ft1* mutants, we used a log2FC ≥ 1 or ≤ -1 for *ft1* mutant in GP-fast background, but in *ft1.1* vs. GP-fast and *ft1.2* vs. GP-fast and no log2FC threshold was set for the comparison of *ft1.3* vs. GP. *FT1*-regulated genes were considered if they were either up- or downregulated in all *ft1* mutants at the same stage.

For additional annotation, a strict one-to-one conversion between BaRTv2 and MorexV3 gene models (Mascher et al., 2021) was created for all BaRTv2 identifiers. First, the longest CDS of each gene model was selected, and the respective datasets were aligned against each other in both directions, using BLASTN (version 2.13.0+), default parameters with “-outfmt 6”. The respective outputs were ordered by seqid, bitscore, evalue, and pident. A one-to-one conversion was reported when there was a reciprocal best hit between a gene model in both directions. If there was no one-to-one conversion for the remaining DEGs, they were manually curated from the remaining best-hit alignments in both directions. In addition, the MorexV3 protein sequences (Hv_Morex.pgsb.Jul2020.aa.fa) were aligned with BLASTP (version 2.13.0+) “-outfmt 6 -max_target_seqs 1” against a local BLASTP database of Araport11 (Araport11_pep_20220914_representative_gene_model, Cheng et al., 2017) and the functional annotations were retrieved from Araport11_GFF3_genes_transposons.current.gff (release 2023-01-02 by TAIR).

Gene ontology (GO) term enrichment was performed using ShinyGO 0.80 (FDR cutoff 0.05) (Ge et al., 2020) on DEGs, separated by tissues and into upregulated and downregulated genes. For this, MorexV3 identifiers were used as input data. Not all BaRTv2 IDs could be annotated with a MorexV3 ID, so the number of genes used for GO term enrichment was reduced by approximately 2-5% as the BaRTv2 annotation is more complete (see Supplemental Tables S1, S2 for detailed numbers). The FDR cut-off for the GO term enrichment was set at 0.05. Top GO terms, including fold enrichment and FDR values, are listed in Supplemental Datasets S3 and S4. Co-expression analysis was conducted using the gene expression of *FT1*-regulated genes in the leaf and MSA, respectively. Co-expression modules were identified using the WGCNA package in R with a signed hybrid network based on biweight midcorrelation (bicor, maxPOutliers=0.05) (Langfelder and Horvath, 2008). The modules were merged for MSA samples using the Dynamic Hybrid Tree Cut algorithm (cutreeDynamic) with minModuleSize=10, cutHeight=0.995. Module “grey” contains the genes that cannot be signed into other modules. The gene lists for each module can be found in Supplemental Datasets S1 and S2. Pearson correlations between module eigengenes and traits were computed, and corresponding *p*-values were estimated using the Student asymptotic test implemented in WGCNA.

### Sugar extraction from leaves and spikes

GP, GP fast, *ft1.1*, *ft1.2*, and *ft1.3* plants were sown in QuickPot 96T trays (HerkuPlast Kubern GmbH, pot volume 75 cm^3^) and transferred to plant growth chambers after stratification and cultivated under LD conditions as described above.

To measure the concentration of soluble sugars and starch in leaves, the youngest, fully elongated leaf (leaf 3 for all genotypes) was sampled 16 DAE.

To represent a full leaf, four small sections (ca. 1 cm) of the tip, base, and mid-section of the leaf were sampled into a 1.5 ml tube and frozen quickly in liquid nitrogen. Plants were sampled at ZT 0 (end of night, EON) and ZT 16 (end of day, EOD). Ten biological replicates were sampled for each genotype and time point. Each biological replicate consisted of leaf material from a single plant.

To measure the concentration of soluble sugars and starch in developing spikes, full spikes were sampled at W8.0-W8.5 and frozen quickly in liquid nitrogen. Plants were sampled at ZT 0 (EON) and 16 (EOD). Ten biological replicates were sampled for each genotype and time point. Each biological replicate consisted of 1-2 spikes.

To extract soluble sugars, leaves were boiled twice in 750 µl 80% EtOH at 85°C for 45 min while shaking (1000 rpm). The supernatant (1.5 ml) was transferred to a fresh tube and concentrated in a vacuum concentrator over night at 35°C and resuspended in 900 µl H2O. The content of glucose, fructose, and sucrose was determined for each sample by subsequently adding 2.5 µg glucose-6-phosphate dehydrogenase (Roche, REF 10127671001), 0.75 U hexokinase (Roche, REF 11426362001), 5 µg glucose-6-phosphate-isomerase (Roche, REF 10128139001), and at least 50 µg invertase (Sigma-Aldrich, Cat. No. I4504) to the samples and measuring the glucose-6-phosphate-dehydrogenase-dependent NADPH accumulation at 340 nm in a microplate reader.

To extract starch, the insoluble fraction (the leaf) was dried for 60 min to remove all EtOH. The leaf was frozen in liquid nitrogen and ground using 2 mm glass beads in a TissueLyser (QIAGEN) and boiled in 800 µl 0.2 M KOH for 45 min at 90°C while shaking (1000 rpm). Samples were neutralized to pH 6.8 with 1 M acetic acid. Starch was digested in each sample by adding 3.6 U alpha-amylase (Sigma Aldrich, Cat. No. 10065-10G) and 2.5 U amyloglucosidase (Sigma Aldrich, Cat. No. 10115-1G-F) at 30°C overnight. The starch amount was measured as glucose content as described above.

### Statistical analyses

All statistical tests were performed using R (RStudio Team, 2022). Significance between two groups in soluble sugars and starch measurement, statistic was determined by comparing GP or *ft1* mutants to GP-fast at either EOD or EON using Fisher’s F-test for homoscedasticity, followed by Welch’s t-statistic. Significance between more than two groups was determined using a one-way ANOVA (function *aov*) and a subsequent Tukey test (function *HSD.test* from package *agricolae*, v1.3-5), *p*-value cutoff at ≤ 0.05. Polynomial regressions (Loess smooth line) were calculated with a 95% confidence interval.

## References

Abe M, Kobayashi Y, Yamamoto S, Daimon Y, Yamaguchi A, Ikeda Y, Ichinoki H, Notaguchi M, Goto K, Araki T (2005) Fd, a bZIP protein mediating signals from the floral pathway integrator FT at the shoot apex. Science (New York, N.Y.), 309(5737): 1052–1056.

Ahn JH, Miller D, Winter VJ, Banfield MJ, Lee JH, Yoo SY, Henz SR, Brady RL, Weigel D (2006) A divergent external loop confers antagonistic activity on floral regulators FT and TFL1. The EMBO Journal, 25(3): 605–614.

Albani MC, Coupland G (2010) Comparative analysis of flowering in annual and perennial plants. Current Topics in Developmental Biology, 91: 323–348.

Babicki S, Arndt D, Marcu A, Liang Y, Grant JR, Maciejewski A, Wishart DS (2016) Heatmapper: Web-enabled heat mapping for all. Nucleic Acids Research, 44(W1): W147–53.

Balanzà V, Martínez-Fernández I, Sato S, Yanofsky MF, Kaufmann K, Angenent GC, Bemer M, Ferrándiz C (2018) Genetic control of meristem arrest and life span in Arabidopsis by a FRUITFULL-APETALA2 pathway. Nature Communications, 9(1): 565.

Banfield MJ, Brady RL (2000) The structure of Antirrhinum centroradialis protein (CEN) suggests a role as a kinase regulator. Journal of Molecular Biology, 297(5): 1159–1170.

Bartlett ME, Thompson B (2014) Meristem identity and phyllotaxis in inflorescence development. Frontiers in Plant Science, 5: 508.

Beales J, Turner A, Griffiths S, Snape JW, Laurie DA (2007) A pseudo-response regulator is misexpressed in the photoperiod insensitive *Ppd-D1a* mutant of wheat (*Triticum aestivum L.*). TAG. Theoretical and Applied Genetics. Theoretische Und Angewandte Genetik, 115(5): 721–733.

Bennett T, Dixon LE (2021) Asymmetric expansions of FT and TFL1 lineages characterize differential evolution of the EuPEBP family in the major angiosperm lineages. BMC Biol 19, 181. 10.1186/s12915-021-01128-8

Benlloch R, Berbel A, Serrano-Mislata A, Madueño F (2007) Floral initiation and inflorescence architecture: A comparative view. Annals of Botany, 100(3): 659–676.

Beydler B, Osadchuk K, Cheng C-L, Manak JR, Irish EE (2016) The Juvenile Phase of Maize Sees Upregulation of Stress-Response Genes and Is Extended by Exogenous Jasmonic Acid. Plant Physiology, 171(4): 2648–2658.

Bi X, van Esse W, Mulki MA, Kirschner G, Zhong J, Simon R, von Korff M (2019) Centroradialis Interacts with *FLOWERING LOCUS T*-Like Genes to Control Floret Development and Grain Number. Plant Physiology, 180(2): 1013–1030.

Biancucci M, Chirivì D, Baldini A, Badenhorst E, Dobetti F, Khahani B, Formentin E, Eguen T, Turck T, Moore JP, Tavakol E, Wenkel S, Schiavo FL, Ezquer I, Brambilla V, Horner D, Chiara M, Perrella G, Betti C, Fornara F (2025) Mutations in HEADING DATE 1 affect transcription and cell wall composition in rice, Plant Physiology, 197 (4), kiaf120, 10.1093/plphys/kiaf120

Bommert P, Whipple C. (2018) Grass inflorescence architecture and meristem determinacy. Semin Cell Dev Biol. 79:37–47. doi: 10.1016/j.semcdb.2017.10.004. Epub 2017 Oct 14. PMID: 29020602.

Buchmann, Haraldsson EB, Schüller R, Rütjes T, Walla AA, von Korff Schmising M, Liu S (2025) bioRxiv 2025.10.31.685778; doi: 10.1101/2025.10.31.685778

Chailakhyan M (1936) New facts in support of the hormonal theory of plant development. (13): 79–83.

Chardon F, Damerval C (2005). Phylogenomic Analysis of the PEBP Gene Family in Cereals. J Mol Evol 61, 579–590). 10.1007/s00239-004-0179-4

Chen Q, Payyavula RS, Chen L, Zhang J, Zhang C, Turgeon R (2018) *Flowering LOCUS T* mRNA is synthesized in specialized companion cells in *Arabidopsis* and Maryland Mammoth tobacco leaf veins. Proceedings of the National Academy of Sciences of the United States of America, 115(11): 2830–2835.

Chen R, Le Luo, Li K, Li Q, Li W, Wang X (2023) Dormancy-Associated Gene 1 (OsDRM1) as an axillary bud dormancy marker: Retarding Plant Development, and Modulating Auxin Response in Rice (Oryza sativa L.). Journal of Plant Physiology, 291: 154117.

Cheng C-Y, Krishnakumar V, Chan AP, Thibaud-Nissen F, Schobel S, Town CD (2017) Araport11: A complete reannotation of the *Arabidopsis thaliana* reference genome. The Plant Journal, 89(4): 789–804.

Corbesier L, Vincent C, Jang S, Fornara F, Fan Q, Searle I, Giakountis A, Farrona S, Gissot L, Turnbull C, et al. (2007) Ft protein movement contributes to long-distance signaling in floral induction of *Arabidopsis*. Science (New York, N.Y.), 316(5827): 1030– 1033.

Coulter M, Entizne JC, Guo W, Bayer M, Wonneberger R, Milne L, Schreiber M, Haaning A, Muehlbauer GJ, McCallum N, et al. (2022) Bartv2: A highly resolved barley reference transcriptome for accurate transcript-specific RNA-seq quantification. The Plant Journal : For Cell and Molecular Biology, 111(4): 1183–1202.

Debernardi JM, Woods DP, Li K, Li C, Dubcovsky J (2022) Mir172-APETALA2-like genes integrate vernalization and plant age to control flowering time in wheat. PLoS Genetics, 18(4): e1010157.

Diallo AO, Agharbaoui Z, Badawi MA, Ali-Benali MA, Moheb A, Houde M, Sarhan F (2014) Transcriptome analysis of an mvp mutant reveals important changes in global gene expression and a role for methyl jasmonate in vernalization and flowering in wheat. Journal of Experimental Botany, 65(9): 2271–2286.

Digel B, Pankin A, von Korff M (2015) Global Transcriptome Profiling of Developing Leaf and Shoot Apices Reveals Distinct Genetic and Environmental Control of Floral Transition and Inflorescence Development in Barley. The Plant Cell, 27(9): 2318–2334.

Digel B, Tavakol E, Verderio G, Tondelli A, Xu X, Cattivelli L, Rossini L, von Korff M (2016) Photoperiod-H1 (*Ppd-H1*) Controls Leaf Size. Plant Physiology, 172(1): 405–415.

Dixon LE, Farré A, Finnegan EJ, Orford S, Griffiths S, Boden SA (2018) Developmental responses of bread wheat to changes in ambient tempera-ture following deletion of a locus that includes FLOWERING LOCUS T1. Plant Cell Environ, 41(7).

Dresselhaus T, Balboni M, Berg L, Dolata A, Hochholdinger F, Huang Y, Jiang G, von Korff M, Ku J-C, van der Linde K, Maika J, Mondragon CL, Raissig MT, Schnittger A, Schnurbusch T, Simon R, Stahl Y, Timmermans M, Thirulogachandar V, Zhao, Yaping Zhou S (2025) How meristems shape plant architecture in cereals—Cereal Stem Cell Systems (CSCS) Consortium, The Plant Cell, 37 (7) koaf150, 10.1093/plcell/koaf150

Ejaz M, von Korff M (2017) The Genetic Control of Reproductive Development under High Ambient Temperature. Plant Physiology, 173(1): 294–306.

Ewels P, Magnusson M, Lundin S, Käller M (2016) Multiqc: Summarize analysis results for multiple tools and samples in a single report. Bioinformatics (Oxford, England), 32(19): 3047–3048.

Faure S, Higgins J, Turner A, Laurie DA (2007) The *FLOWERING LOCUS T*-like gene family in barley (*Hordeum vulgare*). Genetics, 176(1): 599–609.

Fichtner F, Lunn JE (2021) The Role of Trehalose 6-Phosphate (Tre6P) in Plant Metabolism and Development. Annual Review of Plant Biology, 72: 737–760.

Finnegan EJ, Ford B, Wallace X, Pettolino F, Griffin PT, Schmitz RJ, Zhang P, Zhang P, Barrero JM, Hayden MJ, Boden SA, Cavanagh CA, Swain SM, Trevaskis B (2018) Zebularine treatment is associated with deletion of FT-B1 leading to an increase in spikelet number in bread wheat. Plant Cell Environ, 41:1346–1360.

Ge SX, Jung D, Yao R (2020) Shinygo: A graphical gene-set enrichment tool for animals and plants. Bioinformatics (Oxford, England), 36(8): 2628–2629.

Gerasimova SV, AM Korotkova, CW Hertig, S Hiekel, R Hoffie, N Budhagatapalli, I Otto, G Hensel, VK Shumny, AV Kochetov, J Kumlehn, and EK Khlestkina (2018) Targeted genome modification in protoplasts of a highly regenerable Siberian barley cultivar using RNA-guided Cas9 endonuclease. Vavilov Journal of Genetics and Breeding 22, 1033–1039

Gol L, Haraldsson EB, von Korff M (2021) Ppd-H1 integrates drought stress signals to control spike development and flowering time in barley. Journal of Experimental Botany, 72(1): 122–136.

Gol L, Tomé F, von Korff M (2017) Floral transitions in wheat and barley: interactions between photoperiod, abiotic stresses and nutrient status. Journal of Experimental Botany, 68 (7), 1399–1410.

González-Suárez P, Walker CH, Bennett T (2023) Flowering LOCUS T mediates photo-thermal timing of inflorescence meristem arrest in Arabidopsis thaliana. Plant Physiology, 192(3): 2276–2289.

Greenup AG, Sasani S, Oliver SN, Talbot MJ, Dennis ES, Hemming MN, Trevaskis B (2010) *Oddsoc2* is a MADS box floral repressor that is down-regulated by vernalization in temperate cereals. Plant Physiology, 153(3): 1062–1073.

Guo W, Tzioutziou NA, Stephen G, Milne I, Calixto CP, Waugh R, Brown JWS, Zhang R (2021) 3d RNA-seq: A powerful and flexible tool for rapid and accurate differential expression and alternative splicing analysis of RNA-seq data for biologists. RNA Biology, 18(11): 1574–1587.

Halliwell J, Borrill P, Gordon A, Kowalczyk R, Pagano, ML, Saccomanno B, Bentley AR, Uauy C, Cockram J (2016) Systematic investigation of FLOWERING LOCUS T-like Poaceae gene families identifies the short-day expressed flowering pathway gene, TaFT3 in wheat (Triticum aestivum L.). Frontiers in Plant Science, 7, 857.

Hanzawa Y, Money T, Bradley D (2005) A single amino acid converts a repressor to an activator of flowering. Proceedings of the National Academy of Sciences of the United States of America, 102(21): 7748–7753.

Hartmann U, Höhmann S, Nettesheim K, Wisman E, Saedler H, Huijser P (2000) Molecular cloning of SVP: A negative regulator of the floral transition in Arabidopsis. The Plant Journal : For Cell and Molecular Biology, 21(4): 351–360.

John S, Apelt F, Kumar A, Bents D, Annunziata MG, Fichtner F, Mueller-Roeber B, Olas JJ (2022). Transcription factor HSFA7b controls ethylene signaling and meristem maintenance at the shoot apical meristem during thermomemory. 10.1101/2022.10.26.513826

John S., Apelt F., Kumar A., Acosta I.F., Bents D., Annunziata M.G., Fichtner F., Gutjahr C., Mueller-Roeber B., and Olas J.J. (2024). The transcription factor HSFA7b controls thermomemory at the shoot apical meristem by regulating ethylene biosynthesis and signaling in Arabidopsis. Plant Comm. 5, 100743

Jones H, Leigh FJ, Mackay I, Bower MA, Smith LMJ, Charles MP, Jones G, Jones MK, Brown TA, Powell W (2008) Population-based resequencing reveals that the flowering time adaptation of cultivated barley originated east of the Fertile Crescent. Molecular Biology and Evolution, 25(10): 2211–2219.

Kardailsky I, Shukla VK, Ahn JH, Dagenais N, Christensen SK, Nguyen JT, Chory J, Harrison MJ, Weigel D (1999) Activation tagging of the floral inducer *FT*. Science (New York, N.Y.), 286(5446): 1962–1965.

Kumar R, Brar MS, Kunduru B, Ackerman AJ, Yang Y, Luo F, Saski CA, Bridges WC, de Leon N, McMahan C, Kaeppler SM, Sekhon RS (2023) Genetic architecture of source–sink-regulated senescence in maize, Plant Physiology 193 (4): 2459–2479, 10.1093/plphys/kiad460

Lan T, Walla A, Ecem Çolpan Karışan K, Buchmann G, Wewer V, Metzger S, Vardanega I, Haraldsson EB, Helmsorig G, Thirulogachandar V, Simon R, von Korff M (2025) PHOTOPERIOD 1 enhances stress resistance and energy metabolism to promote spike fertility in barley under high ambient temperatures, Plant Physiology, 197(4) kiaf118, 10.1093/plphys/kiaf118

Li C, Lin H, Dubcovsky J (2015a) Factorial combinations of protein interactions generate a multiplicity of florigen activation complexes in wheat and barley. The Plant Journal, 84(1): 70–82.

Li K, Debernardi JM, Li C, Lin H, Zhang C, Jernstedt J, von Korff M, Zhong J, Dubcovsky J (2021) Interactions between SQUAMOSA and SHORT VEGETATIVE PHASE MADS-box proteins regulate meristem transitions during wheat spike development, The Plant Cell, 33(12): 3621–3644, 10.1093/plcell/koab243

Liu W-C, Han T-T, Yuan H-M, Yu Z-D, Zhang L-Y, Zhang B-L, Zhai S, Zheng S-Q, Lu Y-T (2017) Catalase2 functions for seedling postgerminative growth by scavenging H2 O2 and stimulating ACX2/3 activity in Arabidopsis. Plant, Cell & Environment, 40(11): 2720–2728.

Lv B, Nitcher R, Han X, Wang S, Ni F, Li K, Pearce S, Wu J, Dubcovsky J, Fu D (2014) Characterization of *FLOWERING LOCUS T1 (FT1)* gene in Brachypodium and wheat. PloS One, 9(4): e94171.

Lv X, Zhang Y, Zhang Y, Fan S, Kong L.(2020) Source-sink modifications affect leaf senescence and grain mass in wheat as revealed by proteomic analysis. BMC Plant Biol, 20, 257. 10.1186/s12870-020-02447-8

Marthe C, J Kumlehn, and G Hensel (2015) Barley (Hordeum vulgare L.) transformation using immature embryos. In: Wang K (ed.): Agrobacterium Protocols, Volume 1, Methods in Molecular Biology, Vol. 1223, Chapter 6, Springer Science+Business Media: New York, p. 71–83

Mascher M, Wicker T, Jenkins J, Plott C, Lux T, Koh CS, Ens J, Gundlach H, Boston LB, Tulpová Z, et al. (2021) Long-read sequence assembly: A technical evaluation in barley. The Plant Cell, 33(6): 1888–1906.

Melzer S, Lens F, Gennen J, Vanneste S, Rohde A, Beeckman T (2008) Flowering-time genes modulate meristem determinacy and growth form in Arabidopsis thaliana. Nature Genetics, 40(12): 1489–1492.

Miryeganeh M (2021) Plants’ Epigenetic Mechanisms and Abiotic Stress. Genes, 12(8).

Muggeo VMR (2003) Estimating regression models with unknown break-points. Statistics in Medicine, 22(19): 3055–3071.

Muggeo VMR (2008) segmented: An R Package to Fit Regression Models with Broken-Line Relationships. R News. (8/1): 20–25.

Mulki MA, Bi X, von Korff M (2018) *Flowering LOCUS T3* Controls Spikelet Initiation But Not Floral Development. Plant Physiology, 178(3): 1170–1186.

Olas, J.J., Apelt, F., Annunziata, M.G., John, S., Richard, S.I., Gupta, S., Kragler, F., Balazadeh, S., and Mueller-Roeber, B. (2021a). Primary carbohydrate metabolism genes participate in heat-stress memory at the shoot apical meristem of Arabidopsis thaliana. Mol. Plant 14:1508–1524

Ono M, Isono K, Sakata Y, Taji T (2021) Catalase2 plays a crucial role in long-term heat tolerance of Arabidopsis thaliana. Biochemical and Biophysical Research Communications, 534: 747–751.

Matthew J. Paul, Foyer C (2001) Sink regulation of photosynthesis, Journal of Experimental Botany, 52 (360): 1383–1400, 10.1093/jexbot/52.360.1383

Park SJ, Jiang K, Tal L, Yichie Y, Gar O, Zamir D, Eshed Y, Lippman ZB (2014) Optimization of crop productivity in tomato using induced mutations in the florigen pathway. Nature Genetics, 46, 1337–1342.

Patro R, Duggal G, Love MI, Irizarry RA, Kingsford C (2017) Salmon provides fast and bias-aware quantification of transcript expression. Nature Methods, 14(4): 417–419.

Peterson R, Slovin JP, Chen C (2010) A simplified method for differential staining of aborted and non-aborted pollen grains. International Journal of Plant Biology, 1(2): 13.

Pieper R, Tomé F, von Korff M (2020). *FLOWERING LOCUS T4 (HvFT4)* delays flowering and decreases floret fertility in barley. Journal of Experimental Botany 180. 10.1101/2020.03.26.010033

Robinson MD, McCarthy DJ, Smyth GK (2010) Edger: A Bioconductor package for differential expression analysis of digital gene expression data. Bioinformatics (Oxford, England), 26(1): 139–140.

RStudio Team (2022). RStudio: Integrated Development for R. RStudio [Computer software]. Boston, MA: PBC: PBC. Retrieved from http://www.rstudio.com/.

Sang N, Ma B, Liu H, Feng T, Huang X (2025) CRISPR/Cas9-mediated GhFT-targeted mutagenesis prolongs indeterminate growth and alters plant architecture in cotton. Plant Science, 352:112374

Sewelam N, Kazan K, Schenk PM.(2016) Global plant stress signaling: reactive oxygen species at the cross-road. Front Plant Sci:187. 10.3389/fpls.2016.00187

Schindelin J, Arganda-Carreras I, Frise E, Kaynig V, Longair M, Pietzsch T, Preibisch S, Rueden C, Saalfeld S, Schmid B, et al. (2012) Fiji: An open-source platform for biological-image analysis. Nature Methods, 9(7): 676–682.

Shaw LM, Lyu B, Turner R, Li C, Chen F, Han X, Fu D, Dubcovsky J (2019) *Flowering LOCUS T2* regulates spike development and fertility in temperate cereals. Journal of Experimental Botany, 70(1): 193–204.

Shaw LM, Turner AS, Herry L, Griffiths S, Laurie DA (2013) Mutant alleles of *Photoperiod-1* in wheat (*Triticum aestivum L*.) that confer a late flowering phenotype in long days. PloS One, 8(11): e79459.

Shanmugaraj N, Rajaraman J, Kale S, Kamal R, Huang Y, Thirulogachandar V, Budhagatapalli N, Tandron Moya Y A, Hajirezaei M. R., Rutten T, Hensel G, Melzer M, Kumlehn J, Von Wirén N, Mock H, Schnurbusch T (2023). Multilayered regulation of developmentally programmed pre-anthesis tip degeneration of the barley inflorescence. The Plant Cell, 35(11), 3973–4001.

Tamaki S, Matsuo S, Wong HL, Yokoi S, Shimamoto K (2007) Hd3a protein is a mobile flowering signal in rice. Science (New York, N.Y.), 316(5827): 1033–1036.

Tamura K, Stecher G, Kumar S (2021) Mega11: Molecular Evolutionary Genetics Analysis Version 11. Molecular Biology and Evolution, 38(7): 3022–3027.

Taoka K, Ohki I, Tsuji H, Furuita K, Hayashi K, Yanase T, Yamaguchi M, Nakashima C, Purwestri YA, Tamaki S, et al. (2011) 14-3-3 proteins act as intracellular receptors for rice Hd3a florigen. Nature, 476(7360): 332–335.

Thirulogachandar V, Schnurbusch T (2021) ’spikelet stop’ determines the maximum yield potential stage in barley. Journal of Experimental Botany, 72(22): 7743–7753.

Trevaskis B, Tadege M, Hemming MN, Peacock WJ, Dennis ES, Sheldon C (2007) Short Vegetative Phase-Like MADS-Box Genes Inhibit Floral Meristem Identity in Barley. Plant Physiology, 143(1): 225–235.

Turner A, Beales J, Faure S, Dunford RP, Laurie DA (2005) The pseudo-response regulator *Ppd-H1* provides adaptation to photoperiod in barley. Science (New York, N.Y.), 310(5750): 1031–1034.

Waddington SR, Cartwright PM, Wall PC (1983) A Quantitative Scale of Spike Initial and Pistil Development in Barley and Wheat. Annals of Botany, 51(1): 119–130.

Walker CH, Bennett T (2023). FLOWERING LOCUS T mediates photo-thermal timing of inflorescence meristem arrest in Arabidopsis thaliana. Plant Physiology, 192(3), 2276–2289.

Walla A, van Esse WG, Kirschner GK, Guo G, Brünje A, Finkemeier I, Simon R, von Korff M (2020), An Acyl-CoA N-Acyltransferase Regulates Meristem Phase Change and Plant Architecture in Barley, Plant Physiology, Volume 183 (3): 1088–1109, 10.1104/pp.20.00087

Wang B, Smith SM, Li J (2018) Genetic Regulation of Shoot Architecture. Annual Review of Plant Biology, 69: 437–468.

Wang H, Li Y, Pan J, Lou D, Hu Y, Yu D (2017) The bHLH Transcription Factors MYC2, MYC3, and MYC4 Are Required for Jasmonate-Mediated Inhibition of Flowering in Arabidopsis. Molecular Plant, 10(11): 1461–1464.

Wigge PA, Kim MC, Jaeger KE, Busch W, Schmid M, Lohmann JU, Weigel D (2005) Integration of spatial and temporal information during floral induction in *Arabidopsis*. Science (New York, N.Y.), 309(5737): 1056–1059.

Yan L, Fu D, Li C, Blechl A, Tranquilli G, Bonafede M, Sanchez A, Valarik M, Yasuda S, Dubcovsky J (2006) The wheat and barley vernalization gene *VRN3* is an orthologue of *FT*. Proceedings of the National Academy of Sciences of the United States of America, 103(51): 19581–19586.

Zadoks JC, Chang TT, Konzak CF (1974) A decimal code for the growth stages of cereals. Weed Research, 14(6): 415–421.

Zhou L, Cheung M-Y, Li M-W, Fu Y, Sun Z, Sun S-M, Lam H-M (2010) Rice hypersensitive induced reaction protein 1 (OsHIR1) associates with plasma membrane and triggers hypersensitive cell death. BMC Plant Biology, 10: 290.

